# MutSimABC: A Simulation-Based Approximate Bayesian Computational Framework for Mutation Rate Inference in Long-Lived Trees

**DOI:** 10.64898/2025.12.26.696594

**Authors:** Andrea-Maria Grecu, Allen Rodrigo, Teng Li

## Abstract

Long-lived trees accumulate somatic mutations over centuries, forming genetic mosaics in which branches carry distinct genotypes shaped by meristem development. Elongation dynamics regulate stem cell lineage maintenance, while branching events redistribute mutations, producing genetic patterns that often diverge from the physical tree topology. Tomimoto and Satake (2023) formalized these processes through mechanistic simulations; however, their framework was designed for forward prediction rather than parameter inference. Estimating mutation rates and developmental parameters from observed data remains computationally intractable for likelihood-based methods. We present MutSimABC, an Approximate Bayesian Computation (ABC) framework that extends the Tomimoto & Satake model to enable simulation-based parameter inference. MutSimABC jointly estimates the mutation rate (*μ*), elongation parameter (*StD*), and branching bias (*σ*) by comparing observed and simulated mutation distributions without requiring explicit likelihood functions. Validation across 169 simulated datasets with known parameters achieved complete recovery of mutation rate and branching bias, and 99.4% recovery for elongation parameters, within 95% highest posterior density (HPD) intervals. Applied to genomic sequencing data from *Eucalyptus melliodora*, MutSimABC estimated somatic mutation rates ranging from 2.3 × 10^−10^ to 1 × 10^−10^ per site per year and inferred partially stochastic meristem dynamics. This framework enables joint inference of mutation and developmental parameters, advancing the mechanistic analysis of somatic evolution in plants, with flexibility that extends to any long-lived organism.

## I. INTRODUCTION

Long-lived trees are genetic mosaics: somatic mutations accumulating over centuries create branches with distinct genotypes within a single organism. In *Eucalyptus melliodora*, this mosaicism manifests as branch-specific herbivore resistance, where two of eight major branches developed chemical defences against beetle defoliation through somatic mutations [1], [2]. At broader scales, these mutations maintain genetic diversity in clonal plant populations, influence crop improvement through novel variants, and may drive evolutionary processes including speciation [3], [4]. Accurate estimation of somatic mutation rates is therefore essential for understanding plant evolution, optimizing breeding programs in forestry and agriculture, and managing genetic resources in conservation [5], [6].

Mutation rate estimation in long-lived trees proves uniquely challenging because of the developmental architecture underlying this mosaicism. Unlike animals where germline cells are sequestered early in development, plants maintain somatic cells with pluripotent capacity throughout their lifespans [3]. Mutations occurring in shoot apical meristems (SAMs) propagate through subsequent cell divisions to entire branches and eventually to gametes—meaning somatic mutation patterns directly shape heritable genetic variation. This propagation depends on two developmental processes that vary systematically across plant taxa: elongation (vertical stem growth) and branching (lateral meristem formation) [7].

During elongation, SAMs exhibit species-specific organizational patterns. In structured meristems—common in angiosperms—strict tunica-corpus layering preserves cell lineages [8]. In stochastic meristems—more typical of gymnosperms—periclinal divisions disrupt layer boundaries, causing random sampling of stem cells and consequent mutation loss or fixation through genetic drift (Fig. 1). During branching, a subset of SAM cells contributes to each new axillary meristem. Biased sampling—preferentially selecting cells from specific lineages near the branch initiation site— produces uneven mutation distributions across branches, while unbiased sampling distributes mutations more uniformly (Fig. 1). Critically, these processes need not conform to tree topology: stochastic elongation can create mutation patterns independent of branching structure, and biased branching can generate shared mutations at internal nodes that obscure phylogenetic relationships.

**Fig. 1.**
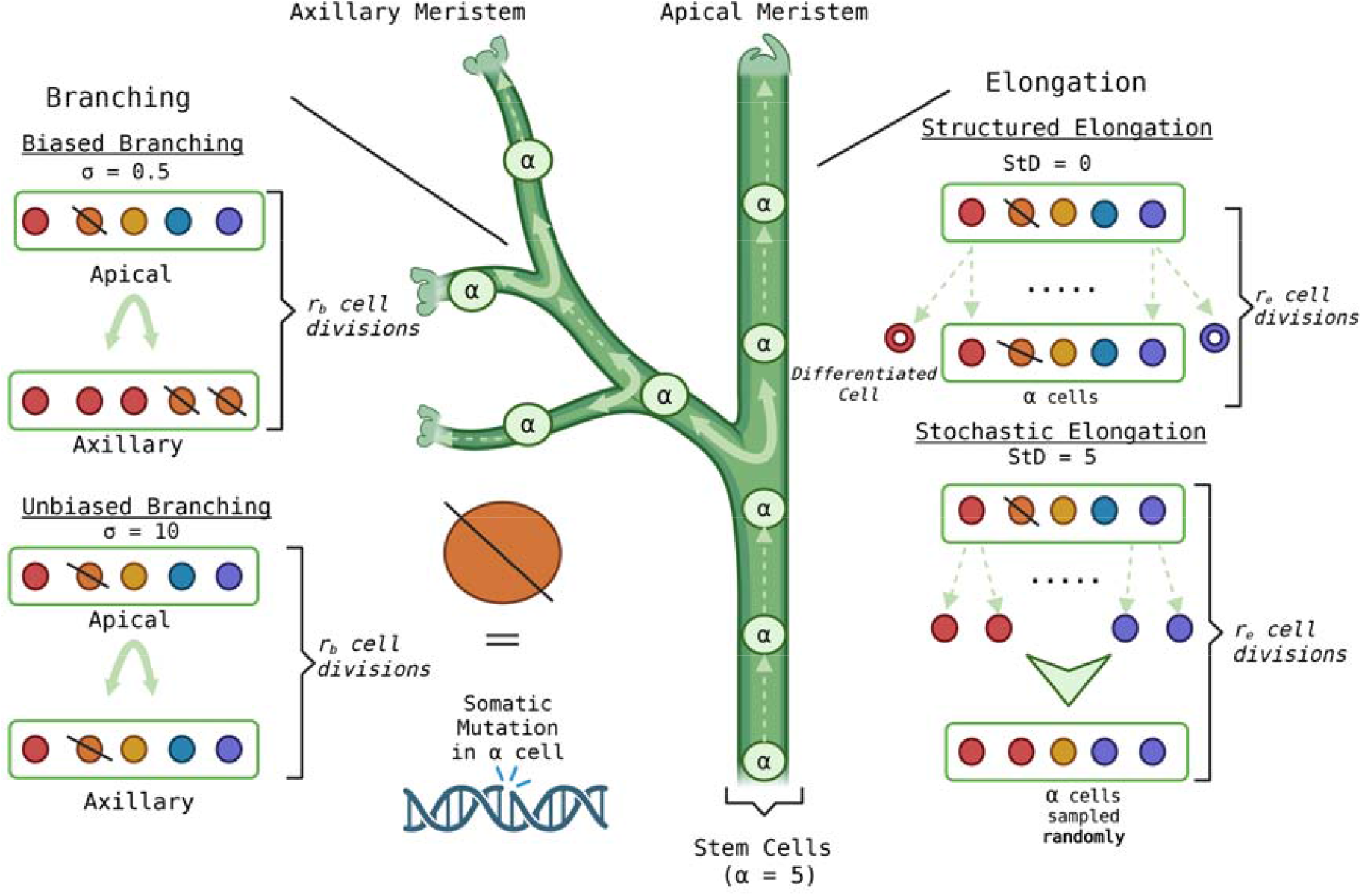
Schematic Representation of Elongation and Branching in Meristems. Meristems (green circles) contain five stem cells (*α* = 5). Dashed arrows represent elongation, with structured elongation (*StD* = 0) preserving lineage and stochastic elongation (*StD* = 5) randomly sampling cell lineages. Solid curved arrows indicate branching, where biased branching (*σ* = 0.5) favors lineage proximity, and unbiased branching (*σ* = 10) ensures equal lineage contribution. Dashed stem cells denote the occurrence of somatic mutations. Parameters *r*_*e*_ = 1 and *r*_*b*_ = 7 represent the number of divisions in elongation and axillary meristem formation, respectively. Created in BioRender. Grecu, A. (2025) https://BioRender.com/rd2vd9c

### A. Current Approaches and Limitations

Whole-genome sequencing (WGS) revolutionized somatic mutation estimation by enabling direct variant detection at nucleotide resolution, consistently revealing lower per-year mutation rates in perennial plants than annuals [9], [10].

However, distinguishing true somatic mutations from sequencing errors and alignment artifacts in organisms where branches carry distinct genotypes remained challenging.

Orr et al. [1] addressed this through a phylogenomic approach using tree topology as a biological filter. By sequencing three biological replicates per branch tip and requiring variants to appear identically across replicates, they eliminated most technical errors [1]. Their method used the physical tree structure as a positive control: phylogenetic trees reconstructed from putative somatic mutations should match the known branching pattern if variants represent true biological signals. Applied to *E. melliodora*, they estimated mutation rates of 1.16 ×10^−10^ to 1.12 × 10^−9^ to per site per year by filtering variants to retain only those whose distribution across branches was consistent with tree topology [1].

This topological filtering assumption has faced critique. Iwasa et al. argued that requiring variants to match branching structure systematically excludes low-frequency mutations, leading to underestimation [11]. More fundamentally, Tomimoto and Satake demonstrated that mutations often do not follow tree topology due to the developmental processes described above [7]. In their analysis of *Populus trichocarpa* [10], they formalized elongation (structured vs. stochastic, parameterized by stem cell lineage maintenance, *StD*) and branching (biased vs. unbiased, parameterized by sampling variance, *σ*) into mechanistic models. Their models incorporating these processes better explained observed mutation distributions than models assuming strict topological congruence, demonstrating that stochastic lineage dynamics and biased sampling create mutation patterns that deviate from physical tree structure [7].

### B. Mechanistic Models and Inference Challenges

Tomimoto and Satake’s framework [7] formalizes elongation (structured vs. stochastic, parameterized by stem cell lineage maintenance, *ScD*) and branching (biased vs. unbiased, parameterized by sampling variance, *σ*). Their models can generate realistic mutation distributions for given parameters but were designed for forward simulation rather than inference. The inverse problem—estimating mutation rate and developmental parameters from observed mutation distributions—is computationally challenging. The stochastic dynamics of meristem elongation, coupled with biased or random sampling during branching and the high dimensionality of tree architectures, make both traditional maximum likelihood and standard Bayesian inference impractical: in both cases, the likelihood is required but cannot be computed.

### C. Approximate Bayesian Computation as a Solution Framework

Approximate Bayesian Computation (ABC) provides a simulation-based, likelihood-free framework ideally suited to this scenario [12]–[14]. Instead of computing *P(data*|*parameters)* directly, ABC samples parameters from prior distributions, simulates data under those parameters, and accepts parameter sets when simulated data sufficiently resemble observed data. ABC has been successfully applied in genomics, epidemiology, and conservation genetics [12]–[16]. Its flexibility allows inference without assuming tree topology, making it ideal for cases where mutations arise through complex meristem dynamics.

### D. Contributions

We introduce MutSimABC, the first inference framework for somatic mutation rates that explicitly accounts for mechanistic meristem dynamics. Our contributions include:

- **Framework Development**: Extension of Tomimoto and Satake’s simulation code into an ABC-Reject pipeline that jointly estimates mutation rate (*μ*), elongation parameter (*μ*), and branching bias (*σ*) from empirical mutation distributions. The framework accommodates diverse input tree topologies and produces posterior distributions with quantified uncertainty via highest posterior density (HPD) intervals.
- **Validation Through Simulation**: Across 169 simulated datasets covering diverse tree architectures, we recovered 100% of mutation rate and branching bias values within 95% HPD intervals and 99.4% of elongation parameter values. Even in cases of high mutational noise or complex topology, effective sample sizes indicated robust posterior exploration.
- **Application to *Eucalyptus melliodora***: Applied to mutation rates of to per empirical genomic data[1],the framework estimated mutation rates of 2.3 ×10^-10^ to 7.1 × 10^-10^ site per year for unfiltered data. The inferred elongation parameter distribution suggested partially stochastic meristem dynamics, providing a biological insight inaccessible to topology-dependent methods.

## II. Methods

### 2. Problem Formulation

We address the inverse problem of estimating somatic mutation rates and meristem developmental parameters from observed mutation distributions in long-lived trees. Given a tree with known topology and branch ages, let D represent observed the distribution of mutations across *n* terminal branches. Our goal is to infer the posterior parameters *θ*={ *μ, ScD, σ*}, where:

- *μ* is the somatic mutation rate per site per year
- *StD* ∈ [0.5] parameterizes elongation dynamics (stem cell lineage maintenance)
- *σ* ∈ [0.5,10] parameterizes branching bias (sampling variance)

The relationship between these parameters and observable mutation patterns is governed by stochastic meristem dynamics (Fig. 1), which vary across tree topologies, precluding the computation of analytical likelihoods and necessitating ABC simulation-inference.

### B. Mechanistic Simulation Model

We extended the forward simulation framework of Tomimoto and Satake [7] to accommodate arbitrary tree topologies and branch numbers. The original implementation was specific to the *Populus trichocarpa* topology studied by Hoffmeister et al. [10]; we generalized the code to accept user-defined tree structures specified via a dictionary-based format that encodes branch ages, connectivity, and hierarchical relationships (see <u>GitHub repository</u> for specifications).

The simulation models somatic mutation accumulation through two coupled developmental processes: **elongation** and **branching** (Fig. 1).

#### Elongation dynamics

During vertical stem growth, *α* = 5 stem cells in the shoot apical meristem (SAM) undergo *r*_*e*_ = 1 cell division per year (Fig. 1, right panel). The parameter quantifies the number of stem cells that do not maintain their lineage:

- When *StD* = 0: structured elongation with perfect lineage preservatWhen *ScD=0* : structured elongation with perfect lineage preservation—one daughter cell differentiates while the other maintains stem cell identity
- When 0 < *StD < 5*: partially stochastic elongation with limited random resampling
- When *ScD* = 5: fully stochastic elongation, where *α* stem cells are randomly sampled from *αr*_*e*_ daughter cells after each division

#### Branching dynamics

Axillary meristem formation involves *r*_*b*_=7 cell divisions producing *α*2*r*_*b*_ candidate cells (Fig. 1, left panel). These cells are spatially arranged on a unit circle, with cells sharing recent ancestry positioned closer together. For each branching event, a position *u* ~ [0,2 *π*) is randomly selected, and *α*=5cells are sampled according to a wrapped normal distribution:

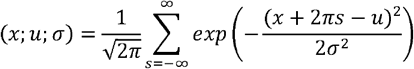

The standard deviation *σ* (biasVar in code) determines sampling bias:

- When *σ* ≥2 *α*: nearly uniform sampling occurs (unbiased branching)—cells are randomly selected regardless of lineage to populate the axillary meristem.
- When 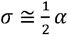 : spatially localized sampling occurs (biased branching)—cells near position *u* preferentially selected, preserving lineage structure.

#### Mutation accumulation

Mutations occur at rate per cell division, each affecting a unique genomic site (Fig. 1, center). The per-site mutation rate is 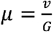 where *G* = 500 Mb. The probability distribution of mutated cells evolves according to:

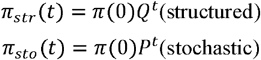

where *Q* and *P* and are transition matrices governing deterministic and stochastic lineage dynamics, respectively [7]. The simulation produces a binary mutation matrix *M ∈*{0,1}^*g*×*n*^indicating the presence or absence of *g* mutations across *n* branches.

#### Implementation details

The simulation pipeline comprises modular functions handling tree topology parsing (`create_tree_list_and_dict`), stem cell mutation tracking (`mutInStemCells`, `mutInBrStemCells`), branching with spatial bias (`sample_mutations`), and mutation matrix construction (`makeMutMatrix`). Core meristem dynamics functions were adapted from Tomimoto and Satake’s original simulation code [7], while topology handling and mutation distribution analysis were developed to enable flexible tree architecture. Complete function descriptions and usage examples are provided in the <u>GitHub repository</u>.

### C. ABC-Reject Inference Framework

Approximate Bayesian Computation (ABC) approximates the posterior distribution *p(θ*/*D)* without ecplicit likelihood evaluation. Our implementation follows the ABC-Reject algorithm:

1. **Prior Specification:** We define uniform priors for our parameterso finterest.

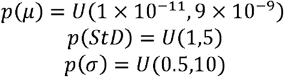

The mutation rate prior was expanded from Orr et al.’s [1] reported range based on preliminary analyses showing posterior density extended beyond initial bounds (Fig. S1, supplementary materials). The *ScD* prior excludes zero because it was found when *StD* = 0, the model reduces to a single-parameter branching-only system, effectively changing the dimensionality of the inference problem. This causes posterior distributions to become trapped at *StD* = 0 with poor mixing across parameter space, a phenomenon known as model-jumping [17]. By restricting to ∈ [1, 5], we maintain consistent model dimensionality while allowing the posterior to identify whether elongation dynamics are partially ( *StD* ≈ 1) or fully ( *StD* ≈ 5) stochastic (*see Supplementary Note S1, supplementary materials*).
2. **Summary Statistics:** For each simulation, we compute branch-specific mutation counts: total mutations per branch, shared mutations between branch pairs, and unique mutations. These statistics capture the essential distributional features while remaining computationally tractable. Let *S*(*D*)denote the summary statistics vector for observed data.
3. **Distance Metric:** We then calculate the Euclidean distance [18] between simulated and observed summary statistics:

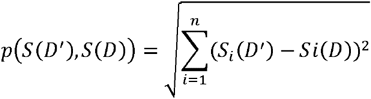

where *n* represents the dimensionality of the vector.
4. **Acceptance Criterion:** Parameter sets, *θ*, are accepted if, *p(S(D*^*′*^)*S*(*D*) ≤ *ε*,where *ε* = 20.This threshold was selected through iterative refinement (aka. sensitivity analysis [19]) to balance posterior informativeness against computational efficiency.
5. **Posterior Approximation:** After *N*=10,000 trials, accepted parameter sets form samples from the approximate posterior *p*(*θ*/*D)*. We computed 95% highest posterior density (HPD) intervals using the ArviZ package [20] to quantify uncertainty.

### D. Validation Strategy

#### Latin Hypercube Sampling for validation design

We generated validation datasets using Latin Hypercube Sampling (LHS) [21], a stratified sampling method that partitions each parameter’s range into equal intervals and samples once from each interval. This ensures comprehensive coverage of multi-dimensional parameter space with fewer samples than grid-based approaches [22], guaranteeing the framework is tested across diverse combinations of tree architectures, mutation rates, and developmental parameters.

#### Dataset generation

We generated 200 synthetic datasets sampling:

- True *μ* values: {6 × 10^−10^,6 × 10^−9^} representing small and large mutation rates
- True *StD* values: {1.25, 2.5, 3.75} (25th, 50th, and 75th percentiles of the prior distribution)
- True *σ* values: {2.875, 5.25, 7.625} (25th, 50th, and 75th percentiles of the prior distribution)
- Tree topologies: 20 unique configurations varying in balance (symmetric vs. asymmetric branching structure) and the ratio of internal to external branch lengths (IE-ratio)

Tree topologies included 4, 6, 8, 10, and 12 terminal branches with combinations of balanced/unbalanced structures and varying IE-ratios. Each tree had a total age ≈ 310 years and a total physical length ≈ 31meters. For each parameter combination, we simulated the “observed” or true known mutation distribution using the mechanistic model (Section II.B). We then applied ABC-Reject with *ε* = 20 (determined via sensitivity analysis on empirical data; Section II.E), accepting 100 parameter sets per validation sample.

#### Evaluation metrics

For each parameter, we assessed coverage [24] (the proportion of validation samples where the true value fell within the 95% highest posterior density (HPD) interval—the narrowest interval containing 95% of posterior probability), effective sample size [25] (ESS, the number of effectively independent samples accounting for autocorrelation, computed via ArviZ [20]), and posterior characteristics including mean, median, credible interval widths, and distance from true values. Of 200 attempted validations, 169 converged within computational constraints (48-hour wall time, 16GB memory per job on NeSI computing cluster). The 169 successful validations provide robust assessment across the majority of biologically relevant parameter space.

### E. Application to E. melliodora Empirical Data

We applied the validated ABC framework to genomic data from *E. melliodora* obtained from Orr et al. [1] (data publicly available at NCBI BioProject PRJNA553104). The input tree topology comprised eight terminal branches with measured physical distances. Branch ages were estimated by converting branch lengths to years using an approximate annual growth rate of 10 cm yr^−1^ derived from *E. grandis* growth studies. Two datasets were analysed:

#### Pre-DNG dataset

330 high-confidence variants identified by standard GATK (Genome Analysis Tool Kit) variant calling [26] without topological filtering. This represents the mutation distribution before phylogenomic quality control.

### Post-DNG dataset

90 variants retained after Orr et al.’s phylogenomic filtering, which excluded mutations inconsistent with tree topology. This represents mutations conforming to the expected phylogenetic structure.

These analyses primarily demonstrate the application of the prototype ABC-Reject framework (MutSimABC) to real biological data. In addition, comparing the pre- and post-filtered datasets provides a means to examine the impact of topological filtering, thereby further contextualising previous critiques of the phylogenomic approach.

#### Sensitivity analysis

Prior to final inference, we conducted iterative sensitivity analyses on both datasets to refine priors *ε* = 19. Initial priors matched Orr et al.’s reported ranges but proved overly restrictive, particularly for the pre-DNG dataset. The mutation rate prior was expanded from [1.6 ×10^-10^,1.12 ×] to 1 ×10^−11^,9 ×10^−9^] based on preliminary posteroior distribution that clustered at prior boundaries. The elongation parameter prior was restricted from [0, 5] to [1, 5] after observing model-jumping artifacts when *StD* = 0 was included. Multiple *ε* values (10, 15, 20, 25, 30) were tested. Lower values (*ε* ≤15) resulted in excessive rejection rates (>99%), while higher values (*ε* ≥ 25) produced posteriors nearly identical to priors. Therefore, we selected *ε* ≥ 25 as the optimal balance, achieving acceptance rates of approximately 3-5% while maintaining informative posteriors distinct from priors.

#### Implementation

For each dataset, we ran 10,000 ABC trials sampling from the refined prior distributions. Each trial simulated mutations across the *E. melliodora* topology using sampled parameters, computed summary statistics, and accepted or rejected based on the distance criterion. Accepted parameter sets were analysed using ArviZ [20] to produce posterior distributions, summary statistics, and 95% HPD intervals for *μ, stD*, and *σ*.

## III. Results

### A. MutSimABC Validation on Simulated Data

We validated MutSimABC on 169 synthetic datasets with known ground-truth parameters spanning diverse tree architectures (4-12 terminal branches, balanced/unbalanced topologies, varying IE-ratios). The framework achieved 100% coverage for mutation rate (*μ*) and branching bias (*σ*), with 99.4% (168/169) coverage for the elongation parameter (*stD*), where coverage is defined as the proportion of datasets in which true values fell within 95% highest posterior density (HPD) intervals.

Representative cases illustrate posterior characteristics across the performance spectrum (Fig. 2, Table I). The best case (unbalanced topology, 8 short branches; Fig. 2A) exhibited tight, well-defined posteriors for all parameters. The mutation rate (*μ*) posterior was unimodal and symmetrical with mean 6.01 × 10^−10^, median 5.91 × 10^−10^ and 95% HPD [2.43 × 10^−10^, 9.44 × 10^−10^], closely bracketing the true value of 6.0 × 10^−10^. The elongation parameter (*StD*) showed slight right skew with mean 2.66, median 2.48, and 95% HPD [1.08, 4.52], successfully capturing the true value of 2.50. Branching bias (*σ*) displayed a broader yet centered posterior with mean 5.12, median 5.03, and 95% HPD [0.57, 9.69], encompassing the true value of 5.25. Effective sample sizes (ESS) were high for developmental parameters (StD ESS=100, *σ* ESS=98), indicating efficient posterior exploration across the 100 accepted samples. The mutation rate exhibited lower ESS (67), likely reflecting its broader posterior distribution and higher sensitivity to data variability, which induces autocorrelation during sampling.

**TABLE I.** VALIDATION POSTERIOR STATISTICS FOR ‘BEST’ AND ‘WORST’ SAMPLES.

|  | Best Sample |  |  | Worst Sample |  |  |
| --- | --- | --- | --- | --- | --- | --- |
| Parameter | Mean (95% HPD) | True Value | ESS | Mean (95% HPD) | True Value | ESS |
| $\mu$ | 6.01 (2.43-9.44)<br>$\times 10^{-10}$ | $6.0 \times 10^{-10}$ | 67 | 3.67 (1.50-6.97)<br>$\times 10^{-10}$ | $6.0 \times 10^{-10}$ | 68 |
| $StD$ | 2.66 (1.08-4.52) | 2.5 | 100 | 2.70 (1.27-4.97) * | 1.25 | 58 |
| $\sigma$ | 5.12 (0.57-9.69) | 5.25 | 98 | 4.95 (0.99-9.54) | 5.25 | 58 |

**Fig. 2.**
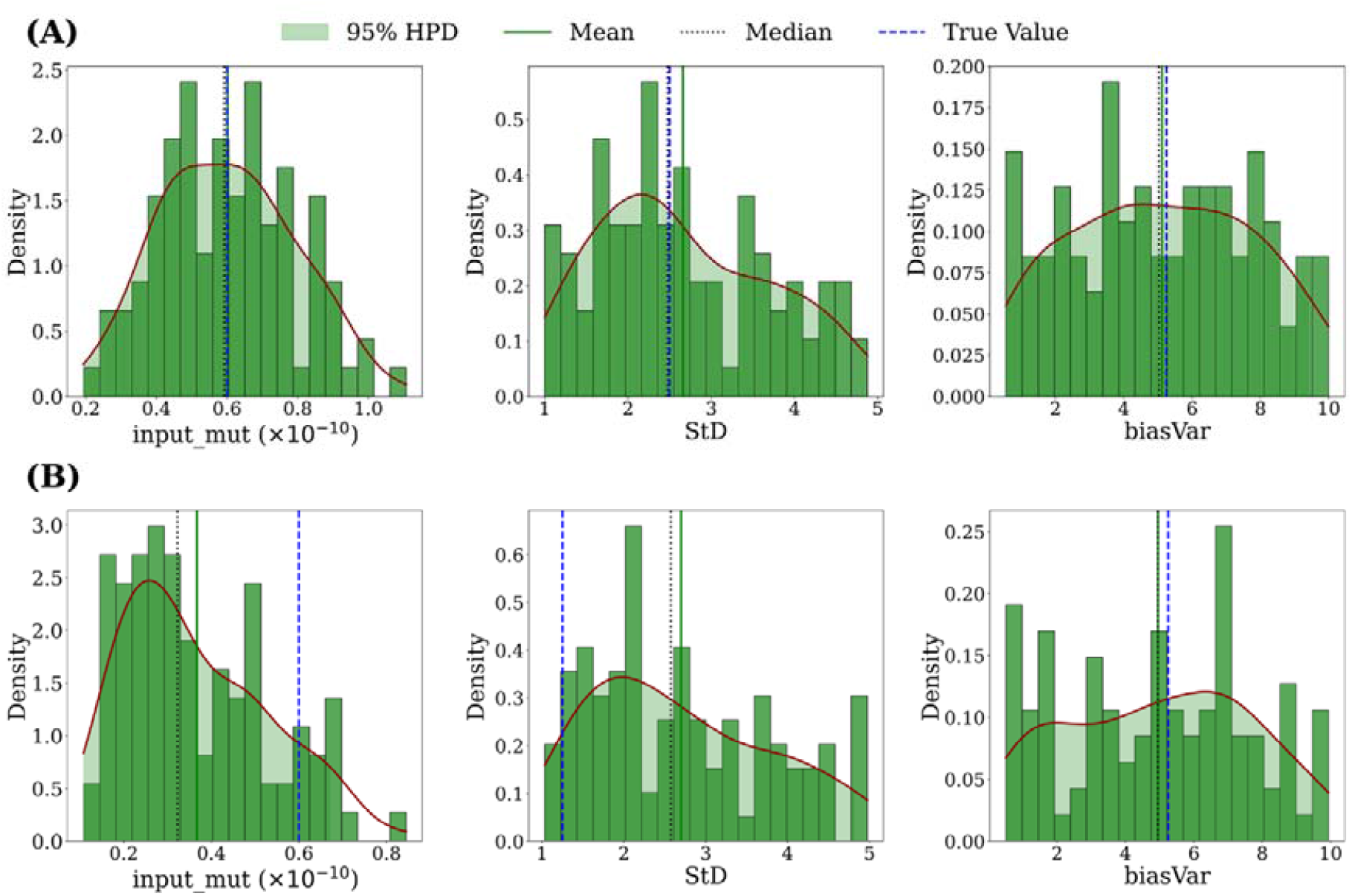
Validation of MutSimABC Framework across Performance Spectrum. Posterior distributions for mutation rate (*input_mut, μ*), elongation parameter (*StD*), and branching bias (*biasVar, σ*) from 169 simulated validation datasets. Green histograms show accepted parameter values; red curves are kernel density estimates; shaded regions indicate 95% highest posterior density (HPD) intervals. Solid green line = posterior mean; dotted green line = median; dashed blue line = true value. **(A)** Representative “best” validation sample (unbalanced topology, 8 short terminal branches) demonstrating optimal recovery with high effective sample sizes (*μ* ESS=67, *StD* ESS=100, *σ* ESS=98). All true values fall within 95% HPD intervals. **(B)** “Worst” validation sample (balanced topology, 8 long terminal branches) representing the single case where *StD* true value (1.25) fell marginally outside 95% HPD [1.27, 4.97] by 0.02 units. Despite this, mutation rate and branching bias were accurately recovered with moderate ESS values (*μ* ESS=68, *StD* ESS=58, *σ* ESS=58), demonstrating framework robustness even under the least favourable conditions.

The single worst case (balanced topology, 8 long branches; Fig. 2B) demonstrated the framework’s performance under challenging conditions. Here, the elongation parameter (*StD*))’s true value (1.25) fell marginally outside the 95% HPD [1.27, 4.97] by 0.02 units—the only such instance among all validations. Despite this, the mutation rate ( *μ*) posterior remained informative, albeit broader than the best case, with mean 3.67 × 10^−10^, median 3.23 × 10^−10^, and 95% HPD [1.50 × 10^−10^, 6.97 × 10^−10^],, successfully capturing the true value of 6.0 × 10^−10^. Branching bias (*σ*) showed moderate precision with mean and median of 4.95 and 95% HPD [0.99, 9.54], including the true value of 5.25. ESS values were lower than the best case (*μ* ESS=68, StD ESS=58, *σ* ESS=58), reflecting reduced mixing efficiency.

Trace plots [26] revealed increased autocorrelation and noisier chains compared to the best case (Supplementary Fig. S2), consistent with the more challenging parameter space and limited number of accepted samples ( *n* = 100). However, the absence of divergent iterations and consistent variability across sampling indicated adequate posterior exploration despite the lower ESS values. The consistency of mutation rate and branching bias recovery across all 169 scenarios, including this least favourable case, confirms the framework’s reliability across diverse tree architectures and parameter combinations.

### B. Appication to Eucalyptus melliodora

We applied MutSimABC to empirical somatic mutation data from *Eucalyptus melliodora* [2], analysing two datasets: 330 high-confidence variants identified before phylogenomic filtering (pre-DNG) and 90 variants retained after topology-based filtering (post-DNG) [1]. Acceptance rates differed substantially between datasets (10.2% for pre-DNG vs. 3.5% for post-DNG), yielding 1,020 and 350 accepted samples, respectively, from 10,000 trials.

#### Mutation rate inference

Pre-DNG analysis produced a unimodal, symmetrical mutation rate (*μ*) posterior (Fig. 3A) with mean 5.3 × 10^−10^, median 5.1 × 10^−10^, and 95% HPD [2.3 × 10^−10^, 7.1 × 10^−10^] per site per year (Table II). This distribution exhibited minimal skew and high bulk ESS (970), indicating efficient posterior exploration. The HPD interval overlapped substantially with Orr et al.’s reported range [1.16 × 10^−10^, 1.12 × 10^−10^] (Fig. 4), despite being derived from unfiltered variants without scaling adjustments.

**Fig. 3.**
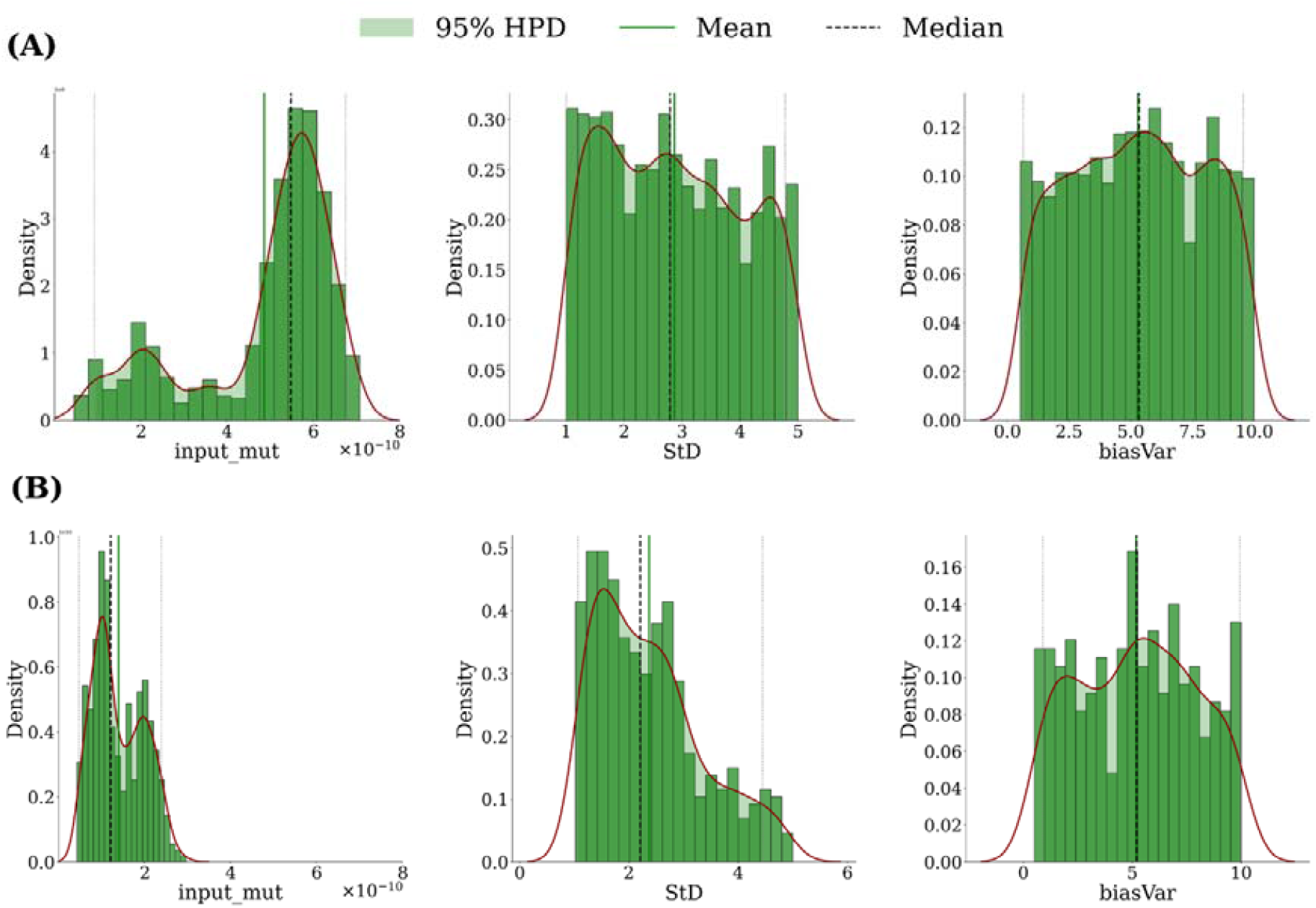
Posterior distributions for *Eucalyptus melliodora* pre-DNG and post-DNG datasets. Mutation rate (*input_mut, μ*, left), elongation parameter (*StD*, centre), and branching bias (*biasVar, σ*, right) posteriors for pre-DNG (panel A, 330 variants) and post-DNG (panel B, 90 variants after topological filtering). Green histograms show accepted parameter values; red curves are kernel density estimates (KDE); shaded regions indicate 95% HPD intervals; solid green line = mean; dashed dark green line = median. Pre-DNG mutation rate displays unimodal, symmetrical distribution. Post-DNG mutation rate exhibits bimodal structure with peaks at approximately 1 × 10^−10^ and 2 × 10^−10^. *StD* posterior shifts from relatively flat (pre-DNG) to skewed toward lower values (post-DNG), indicating structured elongation preference. Branching bias (*σ*) posteriors remain broad and uninformative in both datasets, closely resembling prior distributions. Posterior distributions generated using ArviZ [20].

**Fig. 4.**
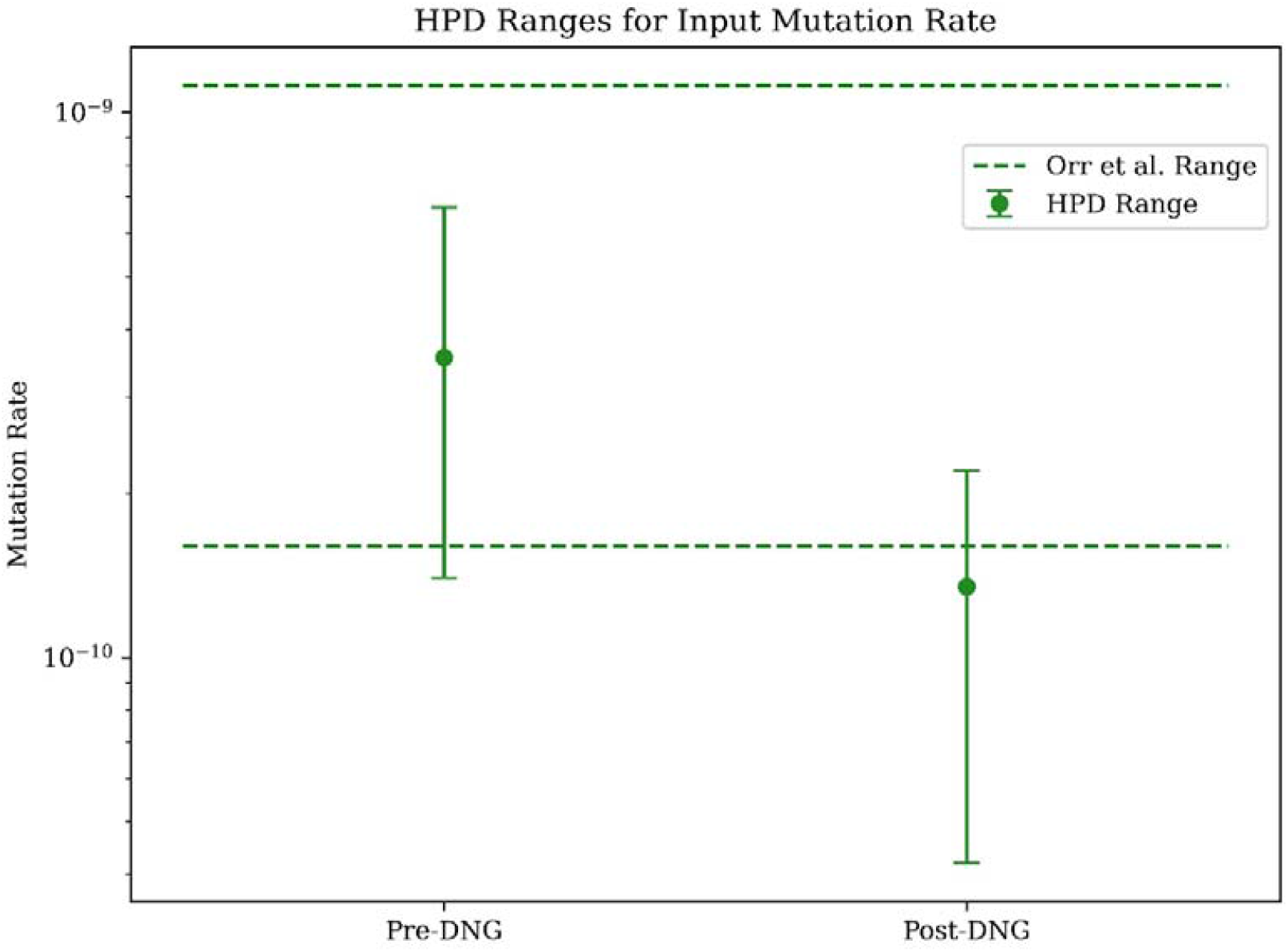
Comparison of mutation rate 95% HPD intervals to previously reported estimates. MutSimABC posterior 95% HPD intervals for pre-DNG (green) and post-DNG (blue) *E. melliodora* datasets compared to Orr et al.’s [1] reported range [1.16 × 10^−10^, 1.12 × 10^−9^] (as noted by the green dashed lines). Pre-DNG HPD [2.3 × 10^−10^, 7.1 × 10^−10^] overlaps substantially with the reported range. Post-DNG HPD [4.2 × 10^−11^, 2.2 × 10^−10^] falls largely below the reported range, with minimal overlap, despite Orr et al.’s scaling adjustments intended to compensate for topological filtering. This discrepancy indicates that phylogenomic filtering removes more mutation signal than previously estimated.

Post-DNG analysis revealed markedly different posterior characteristics (Fig. 3B). The mutation rate (*μ*) distribution was bimodal, with distinct peaks at approximately 1 × 10^−10^, and 2 × 10^−10^, as evidenced by a clear depression in the KDE curve between these regions. The posterior mean was 1.0 × 10^−10^, median 8.5 × 10^−11^, and 95% HPD 4.2 × 10^−11^, 2.2 × 10^−10^, (Table II). This HPD fell largely outside Orr et al.’s reported range (Fig. 4), with minimal overlap at the lower boundary. Bulk ESS (263) and tail ESS (283) were lower than pre-DNG, reflecting the reduced number of accepted samples (350 vs. 1,020) due to the highly filtered nature of the dataset. Trace plots showed relatively consistent variability across iterations despite the shorter chain length (Supplementary Fig. S3), with the lower ESS attributable primarily to branches with reduced mutation signal (i.e., branches 5 and 6 in [1]), where topological filtering retained minimal unique mutations.

**TABLE II.** POSTERIOR STATISTICS FOR *EUCALYPTUS MELLIODORA* DATASETS.

|  | Parameter | Mean | SD | 95% HPD | ESS<br>(Bulk) | ESS<br>Tail |
| --- | --- | --- | --- | --- | --- | --- |
| **Pre-DNG** | $\mu$ | $5.3 \times 10^{-10}$ | $1 \times 10^{-10}$ | $[2.3 \times 10^{-10}, 7.1 \times 10^{-10}]$ | 970 | 707 |
| | $StD$ | 2.943 | 1.174 | [1.002, 4.760] | 914 | 819 |
| | $\sigma$ | 5.307 | 2.681 | [0.639, 9.442] | 842 | 741 |
| **Post-DNG** | $\mu$ | $1.0 \times 10^{-10}$ | $1 \times 10^{-10}$ | $[4.2 \times 10^{-10}, 2.2 \times 10^{-10}]$ | 263 | 283 |
| | $StD$ | 2.36 | 0.980 | [1.033, 4.26] | 342 | 320 |
| | $\sigma$ | 5.18 | 2.730 | [0.991, 9.939] | 282 | 272 |

#### Elongation parameter inference

*StD* posteriors differed between datasets (Fig. 3A, centre). Pre-DNG exhibited a relatively flat, broad distribution with mean 2.94, median 2.85, and 95% HPD [1.00, 4.76] (Table II), indicating limited information to distinguish between partially structured and stochastic elongation modes. Bulk ESS (914) and tail ESS (819) were high [25], with trace plots showing consistent mixing and moderate variability (Supplementary Fig. S3).

Post-DNG analysis revealed a posterior skewed toward lower *StD* values (Fig. 3B, centre), with mean 2.36, median 2.25, and narrower 95% HPD [1.03, 4.26] (Table II). This shift suggests topological filtering preferentially retained mutations consistent with more structured lineage inheritance, where deterministic cell division preserves mutations through defined lineages. ESS values decreased (bulk 342, tail 320), with trace plots showing slower mixing and higher autocorrelation compared to pre-DNG (Supplementary Fig. S3). The appearance of value clustering in trace plots indicated successive samples were more dependent, reducing sampling efficiency.

#### Branching bias inference

Branching bias (*σ*) posteriors were broad and uninformative for both datasets (Fig. 3, right). Pre-DNG showed mean 5.31, median 5.20, and 95% HPD [0.64, 9.44]; post-DNG showed mean 5.18, median 5.15, and 95% HPD [0.99, 9.94] (Table II). Both distributions closely mirrored the uniform prior U(0.5, 10), with means near the prior centre and HPD intervals spanning nearly the entire prior range. Trace plots revealed moderate mixing for pre-DNG (bulk ESS 842, tail ESS 741) and slower mixing for post-DNG (bulk ESS 282, tail ESS 272), with increased clustering evident in the latter (Supplementary Fig. S3). The flat posterior shapes indicate that branching dynamics contributed minimal information to mutation distribution patterns in this system.

## IV. Discussion

### A. Framework Validation and Performance

MutSimABC was validated across 169 simulated datasets with known ground-truth parameters, successfully recovering true values within 95% HPD intervals for all three parameters in nearly all cases: 100% for mutation rate (*μ*), and branching bias (*σ*) and 99.4% for elongation parameter (*StD*). This near-perfect recovery held across diverse tree architectures including 4-12 terminal branches, balanced and unbalanced topologies, and varying internal-to-terminal branch length ratios. Representative cases (Fig. 2, Table I) illustrate posterior characteristics ranging from tight, well-defined distributions to broader yet informative distributions. These validation results establish that the ABC-Reject framework reliably approximates posterior distributions when applied to data generated from the underlying mechanistic model, providing proof-of-concept for the inference approach.

### B. Biological Insights from Empirical Applications

Application of MutSimABC to *E. melliodora* revealed striking differences between pre-DNG (330 variants) and post-DNG (90 variants remaining after topological filtering), providing insights into both mutation rate estimation and the impact of topology-based filtering on inference.

The pre-DNG analysis produced a mutation-rate posterior with a mean(5 × 10^−10^) and 95% HPD [2.30 × 10^−10^, 7.10 × 10^−10^] that substantially overlapped with Orr et al.’s [1] reported range [1.16 × 10^−10^, 1.12 × 10^−9^], despite deriving from unfiltered variants without scaling adjustments (Fig. 4). This concordance validates both MutSimABC’s inference capability and the rationale of pre-phylogenomic-filtering mutation detection. Post-DNG analysis yielded a markedly lower posterior distribution (mean: 1.0 × 10^−10^, 95% HPD: [4.2 × 10^−11^, 2.2 × 10^−10^,]) that fell largely outside Orr et al.’s [1] reported range (Fig. 4). This disparity indicates that phylogenomic or topology-based filtering removed more mutation signal than accounted for by Orr et al.’s scaling adjustments. Topological filtering, by construction, excludes variants inconsistent with tree structure. These are precisely the variants meristem stochasticity and branching bias generate as described in Tomimoto & Satake’s mechanistic model [7]. MutSimABC’s direct inference from observed distributions, without topology-conforming requirements, reveals the true extent of this information loss, reinforcing critiques by Iwasa et al. and Tomimoto & Satake regarding systematic underestimation in topology dependent approaches [7], [11].

The *StD* posterior distributions provide biological insights inaccessible to phylogenomic or other mutation estimation methods. The Pre-DNG dataset returned a relatively flat posterior (mean: 2.94, 95% HPD [1.00, 4.76], Fig. 3A), suggesting the 330 variants provided insufficient signal to definitively resolve elongation behavior. This broad distribution could indicate *E. melliodora* exhibits mixed elongation dynamics or that greater mutation signal would be required to further distinguish stochasticity levels. Analysis of the post-DNG dataset revealed a posterior skewed toward lower *StD* values (mean: 2.36, 95% HPD [1.04, 4.26], Fig. 3B), suggesting that topological filtering preferentially retained mutations consistent with structured lineage inheritance. In structured meristems (*StD* approaching 0), strict cell lineage preservation fixes mutations within defined clades, producing distributions that align with branching topology. As meristems become more stochastic (*StD* approaching 5), random lineage sampling creates mutation patterns that deviate from the physical tree structure. Consequently, topology-based filtering systematically excludes stochastic-elongation signatures, biasing the retained mutation distribution toward structured modes. This interpretation carries an important caveat: the current framework excludes *StD* = 0 due to model-jumping artifacts [17] (discussed in Section IV-C1). If fully structured elongation provides the optimal fit for post-DNG data, the observed skew toward lower *StD* values may represent a boundary effect where the posterior attempts to approach an excluded parameter region [27].

Branching bias (*σ*) posteriors remained broad and uninformative for both datasets, closely mirroring the uniform prior U(0.5, 10) (Fig. 3). Mean values near the prior center (5.31 for pre-DNG, 5.18 for post-DNG) and 95% HPD intervals spanning nearly the entire prior range indicate that branching dynamics contributed minimal information to mutation distribution patterns. Whether this reflects true biological insignificance or model limitations remains unclear. Tomimoto and Satake similarly observed elongation-dominated effects in their analysis of *Populus trichocarpa*, suggesting that axillary meristem formation may play a limited role in shaping mutation accumulation patterns [7], [10]. However, an alternative explanation is that the wrapped normal distribution used to parameterize spatial sampling of stem cells during branch formation may not adequately represent the biological mechanisms of axillary meristem initiation. Testing alternative mathematical representations of lineage sampling—such as models based on cell ancestry, developmental signals, or empirical branching patterns—could clarify whether branching truly has minimal impact or whether current parameterization simply fails to capture relevant biological processes [28].

### C. Methodological Limitations and Challenges

#### C.1 Model-Jumping and Dimensionality Constraints

The observed discontinuity in mutation rate estimates between *StD* = 0 and *StD* ≥ 1 (Supplementary Fig. S1) reveals a fundamental limitation of the ABC-Reject framework. When *StD* = 0, elongation becomes fully structured with deterministic cell division preserving all lineages, eliminating elongation as a source of stochastic variations. Branching bias (*σ*) becomes the sole determinant of mutation distributions, effectively transitioning the model from a 3-parameter space {*μ, StD, σ*} to a 2-parameter space {*μ, σ*}. This creates a sharp boundary rather than continuous parameter variation. In Bayesian inference, this phenomenon is defined as model-jumping, where models with different dimensionalities occupy non-overlapping parameter regions [17]. Standard ABC-Reject assumes continuous parameter spaces and cannot accommodate such discrete shifts [12]-[14].

We restricted the prior range of *StD* to [1,5] to maintain consistent dimensionality and prevent sampling instability. However, this exclusion sacrifices biological realism: highly structured elongation (*StD* ∈[0,1)) represents a plausible developmental mode, particularly in angiosperms with strict tunica-corpus organization [29]. The post-DNG posterior’s skew towards lower *StD* values suggests that fully structured elongation may provide superior fit, yet MutSimABC as it stands cannot explore this hypothesis. Future work could investigate fixing *StD* = 0 as a discrete model alternative, rather than attempting continuous sampling [30], [31]. We would then expect that fully structured elongation would yield even lower mutation rate estimates for the topologically filtered data (post-DNG). Alternatively, Adaptive Sequential Monte Carlo ABC, in which parameters are propagated through progressively stricter tolerance thresholds, could enable smoother transitions between distinct parameter regimes and overcome model-jumping artifacts [32]. While trans-dimensional ABC methods or Reversible Jump MCMC could accommodate variable-dimension inference, their computational expense currently limits practical implementation [33] – [35].

#### C2. Mutation Assignment at Ultra-Low Rates

Tomimoto and Satake’s simulation framework assigns mutations probabilistically [7]: a random number is drawn uniformly from [0, 1] and compared to the mutation rate. A mutation only occurs when this random number falls below the rate itself. At extremely low mutation rates (i.e., < 9.0 × 10^−12^), this criterion becomes so restrictive that mutations are rarely assigned, effectively producing zero-mutation simulations that disrupt downstream functions. While the framework operates reliably across plausible somatic mutation rates for long-lived trees (10^−9^ to 10^−11^), its failure below this range may artificially compress the post-DNG posterior’s lower tail. This limitation could contribute to the observed bimodal structure by forcing accepted parameter sets into higher mutation rate regions when lower rates cannot generate sufficient mutations to match observed data (Fig. 3B). Refinements to mutation assignment strategies—such as Poisson-process implementations [36] or cumulative probability approaches [37]—could enable accurate sampling of ultra-low mutation rates and reveal whether the post-DNG posterior extends further into lower parameter space than currently observed.

#### C.3 Posterior Bimodality and Parameter Interactions

The post-DNG mutation rate posterior exhibited bimodal structure with peaks at approximately 1 × 10^−10^ and 2 × 10^−10^ (Fig. 3B), contrasting sharply with the pre-DNG unimodal distribution. Bimodal posteriors in Bayesian frameworks indicate two competing parameter regions with substantial probability, suggesting the data supports multiple plausible mutation rate estimates rather than a single dominant value [38]. Scatter plot analysis (Supplementary Fig. S1) revealed a negative correlation between mutation rate (*μ*) and *StD*: mutation rates peak when *StD* is low (1-2) and systematically decrease as *StD* increases, clustering into two broad groups. This interaction effect likely drives the observed bimodality.

Topology-based filtering reinforces this relationship by preferentially retaining mutations consistent with tree structure. Structured elongation (low *StD*) fixes more mutations along defined lineages, creating distributions that align with branching topology. Higher mutation rates compensate for structured meristems’ tendency to preserve lineages, producing overall mutation counts consistent with observed data. Conversely, stochastic elongation (higher *StD*) requires lower mutation rates to match filtered distributions, as random lineage sampling already reduces mutation fixation. The exclusion of *StD* = 0 may partially obscure this effect—a fully structured meristem would maximize lineage preservation, potentially creating an even more distinct parameter cluster. Additionally, the mutation assignment limitation (Section IV-C2) could force the higher-rate peak by preventing ultra-low rates from generating sufficient mutations. Together, these factors suggest the bimodal posterior represents topology filtering’s interaction with elongation dynamics, amplified by prior constraints and model limitations. Future work incorporating *StD* = 0 as a discrete alternative and improving ultra-low-rate assignment could clarify whether bimodality persists or resolves into a continuous gradient.

#### C.4 Data Quality Dependencies

Data quality represents a critical determinant of inference success in empirical applications. While validation (Section IV-A) established algorithmic capability under ideal conditions, real-world applications face additional challenges in mutation detection, age estimation, and topological accuracy. Although physical tree topology can be measured directly, converting branch lengths to chronological age introduces substantial uncertainty. This study assumed 10 cm yr□^1^ growth rate based on *E. grandis* studies [39], but growth rates vary with environmental conditions, tree age, and branch position. Inaccurate age estimates systematically bias mutation rate inference, as mutation rate (*μ*) is calculated per site per year. Refining age estimation techniques—through dendrochronology, growth ring analysis, or species-specific allometric models—would directly improve mutation rate accuracy [40].

Empirical data quality critically affects inference outcomes [41]. Validation datasets contain pure mutation signal arising solely from known parameter combinations, enabling the tight, well-defined posteriors observed in Fig. 2A. Empirical data introduces multiple uncertainty sources including: (1) false positives and negatives from variant calling algorithms, (2) variable sequencing coverage across branches, (3) spurious variants from alignment artifacts, and (4) systematic removal of biologically relevant signal through topology-based filtering [42]. These factors reduce information content available for parameter estimation, manifesting as broader credible intervals and less definitive posterior shapes in empirical applications (Fig. 3) despite larger sample sizes (1,020 for pre-DNG, 350 for post-DNG versus 100 for validation). The observed differences between pre-DNG and post-DNG posteriors demonstrate how filtering affects inference, but the quality of the initial 330 variants equally determines whether meaningful biological signal exists to infer.

These data quality dependencies highlight a fundamental limitation: no inference method can extract reliable parameters from low-quality input data. MutSimABC requires accurate mutation calls, minimal filtering artifacts, and precise branch age estimates to produce biologically meaningful posteriors. Future applications should prioritize experimental design optimization—improving variant calling pipelines, optimizing sequencing depth across branches, and refining age estimation methods—alongside computational method refinement.

#### C.5 Computational Efficiency

ABC-Reject’s primary limitation is computational inefficiency. The algorithm samples parameters from priors, simulates data under those parameters, and accepts parameter sets only when Euclidean distance between simulated and observed summary statistics falls below the threshold *ε*. With *ε* = 20, acceptance rates were 10.2% (pre-DNG) and 3.5% (post-DNG), requiring 10,000 trials to obtain 1,020 and 350 accepted samples, respectively. Each trial involves full mutation simulation across the tree topology, making the approach computationally expensive as parameter dimensionality or tree complexity increases.

Sequential Monte Carlo ABC (ABC-SMC) offers potential efficiency gains. ABC-SMC iteratively refines posterior distributions across multiple generations, using adaptive thresholds that progressively tighten tolerance while focusing sampling on high-probability regions [32]. This approach reduces trial counts while maintaining or improving accuracy. ABC-SMC’s iterative framework could additionally address model-jumping by enabling gradual transitions between parameter regimes as tolerance tightens [43]. Implementation of ABC-SMC represents a logical next step for scaling MutSimABC to larger datasets and more complex tree architectures.

### D. Framework Contributions and Future Directions

Despite identified limitations, MutSimABC represents a significant methodological advance for somatic mutation rate estimation in long-lived organisms. As a simulation-based, likelihood-free method, it accommodates complex stochastic processes where analytical likelihood computation is intractable. The framework explicitly incorporates meristem developmental biology through mechanistic models of elongation and branching, providing a biologically informed alternative to methods that assume mutations strictly follow tree topology. By jointly estimating *μ, StD*, and *σ* with quantified uncertainty via HPD intervals, MutSimABC enables hypothesis testing about developmental processes alongside mutation rate inference—capabilities unavailable to existing approaches. The validation success—recovering true parameters across 169 diverse scenarios—establishes proof-of-concept that simulation-based inference can bridge the gap between forward models (which predict observable patterns from parameters) and the inverse problem (estimating parameters from observations).

The framework’s flexibility extends to any long-lived organism with measurable topology and estimable branch ages, provided variant calling achieves sufficient quality. This generality makes MutSimABC particularly valuable where phylogenomic assumptions appear questionable, such as organisms with known stochastic meristem dynamics, species exhibiting adventitious branching, or systems where topological filtering excludes substantial mutation signal.

Immediate refinements should address identified limitations. Implementing ABC-SMC would improve computational efficiency while enabling *StD* = 0 incorporation through trans-dimensional sampling or discrete model comparison. Enhancing mutation assignment algorithms to handle ultra-low rates (< 9.0 × 10^−12^) would allow complete posterior exploration at lower boundaries. Systematic prior sensitivity analysis using perturbation functions and influence diagnostics could formalize the current iterative prior refinement process [44] – [45]. Testing alternative mathematical representations of branching bias might reveal whether current uninformative σ posteriors reflect true biological insignificance or model limitations.

Longer-term extensions should expand biological realism and applicability. Applying MutSimABC to diverse plant species with varying meristem architectures (strict versus plastic, determinate versus indeterminate) would test framework generality and potentially reveal taxon-specific patterns in developmental parameters. Multi-tree inference, pooling mutation data across multiple individuals of the same species, could improve parameter estimation precision by increasing statistical power while simultaneously quantifying individual-level variation around species-level parameter estimates [46]. Integration with improved experimental pipelines including: (1) developing species-specific guidelines for branch age estimation, (2) optimizing variant calling parameters for somatic mutation detection, and (3) designing sequencing strategies that maximize mutation signal—would enhance input data quality. As sequencing costs decline and long-read technologies improve, combining MutSimABC with whole-genome assemblies from multiple branches could enable unprecedented resolution of somatic mutation dynamics. The framework’s current implementation as a prototype demonstrates feasibility while highlighting paths toward a robust, production-ready tool for studying mutation accumulation in organisms where developmental complexity renders traditional methods inadequate.

## Acknowledgments

We thank Sou Tomimoto and Professor Akiko Satake (Mathematical Biology Laboratory, Kyushu University) for generously sharing their simulation code and for their valuable guidance during its implementation. Their support substantially strengthened this work. We acknowledge the use of New Zealand eScience Infrastructure (NeSI) high performance computing facilities. We also acknowledge the use of OpenAI’s ChatGPT-5 & Anthropic’s Claude for assistance in structuring sections of the text, refining written expression, and facilitating idea development, particularly in relation to coding and methodological organization. All research design, intellectual content, methodological decisions, experimental implementations, analyses, and interpretations presented in this publication are solely our own.

## Notes

This work was supported in part by the start-up funds from the University of Auckland, New Zealand under Grant 4020-12090.

### Competing Interest Statement

The authors have declared no competing interest.

