## supplementary materials for "MutSimABC: A Simulation-Based Approximate Bayesian Computational Framework for Mutation Rate Inference in Long-Lived Trees"

**Supplementary Note. S1. Model-Jumping and Dimensionality Reduction**

In the MutSimABC framework, the elongation parameter $\left( StD \right)$ determines stem cell lineage maintenance during meristem growth. When $StD=0$, elongation is fully structured with deterministic cell division preserving all lineages. Critically, this eliminates elongation as a source of stochastic variation, making the branching bias parameter (*σ*) the sole determinant of mutation distribution patterns. This effectively transforms the model from a 3-parameter space {*μ,* $StD$*, σ*} to a 2-parameter space {*μ, σ*}, causing a sharp transition in posterior values at the $StD=0$ boundary.

Such abrupt shifts are characteristic of model-jumping, a well-documented challenge in Bayesian inference where models with different dimensionalities occupy non-overlapping regions of parameter space [17]. Standard ABC-Reject cannot handle these transitions smoothly, as it assumes continuous parameter spaces. Specialized techniques such as Reversible Jump MCMC or trans-dimensional ABC methods are typically required [33]-[35].

To avoid this instability while preserving biological relevance, we restricted the prior to $StD\in[1,5]$ representing a continuous spectrum from partially structured ($StD\approx1$) to fully stochastic ($StD=5$) elongation. Future implementations incorporating adaptive sequential Monte Carlo ABC [32] or hierarchical model selection frameworks could accommodate $StD=0$ as a discrete model alternative, enabling formal comparison of fully structured versus stochastic elongation hypotheses. However, this would necessitate far greater computational resources [34].

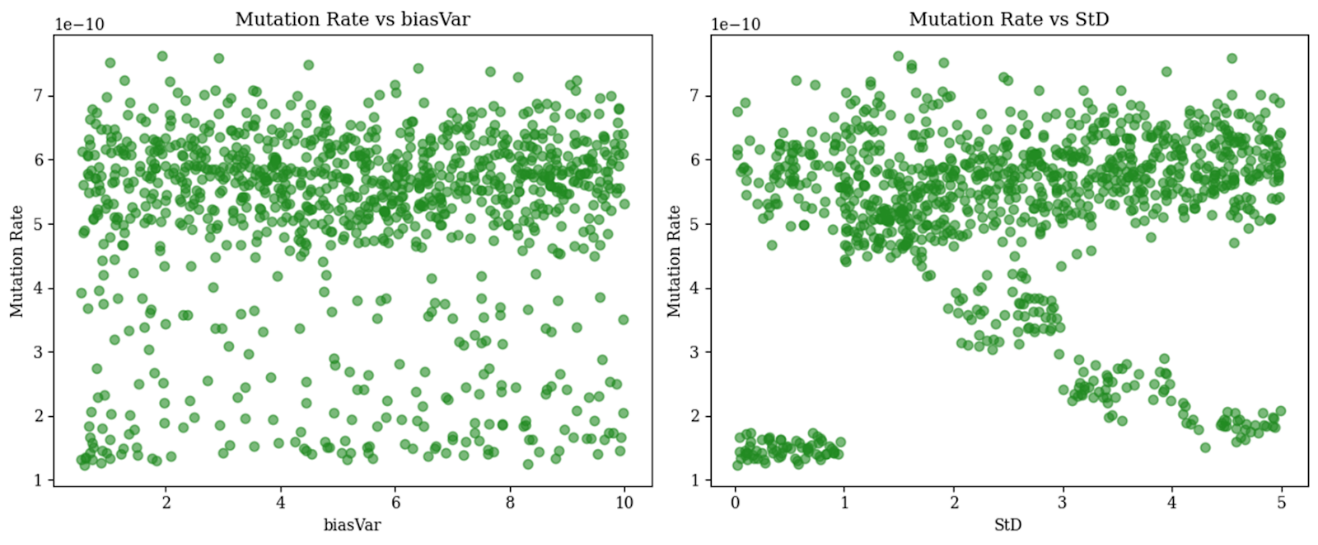

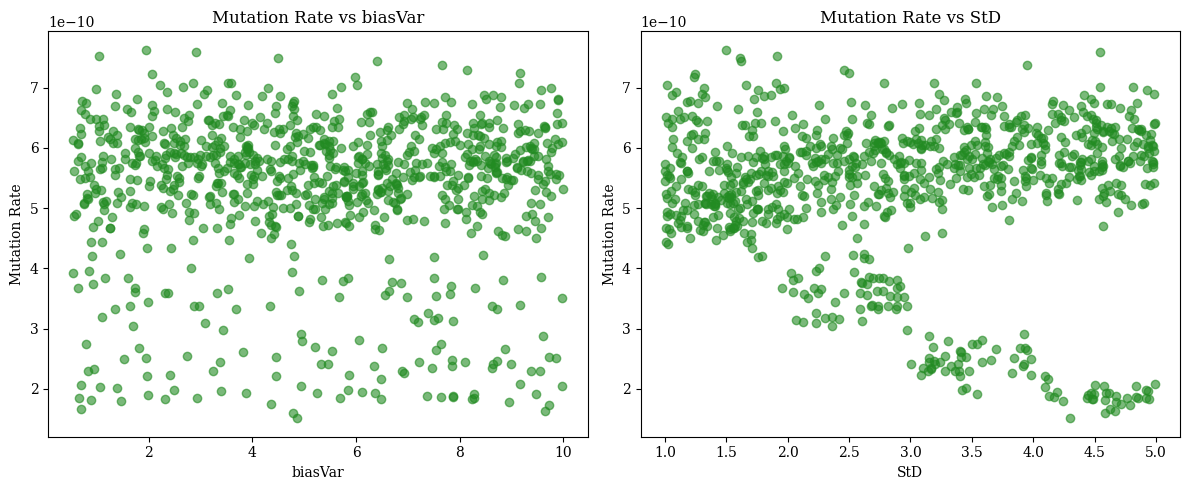

**Supplementary Fig. S1. Model-Jumping Arising from** $\boldsymbol{StD}$**=0 Inclusion.** Scatter plots showing accepted posterior mutation rate ($\mu)$ vs. elongation parameter ($StD$) values for the pre-DNG *E. melliodora* dataset. (Left) When $StD=0$ is included in the prior, mutation rates cluster tightly at lower values with abrupt discontinuity at the $StD=0$ to $StD>0$ transition. (Right) After excluding $StD=0$ from prior, mutation rates distribute more smoothly across $StD\in[1,5]$. This discontinuity arises because $StD=0$ represents a structurally different model where only branching bias ($\sigma$) influences mutation accumulation, effectively reducing parameter dimensionality. This model-jumping behavior, characteristic of Bayesian inference when different models occupy non-overlapping parameter spaces, motivated restricting the prior to $StD\in[1,5]$ to maintain consistent dimensionality throughout inference*.

*A downward trend can be observed in mutation rate estimates across both scatter plots, before and after removing $StD =0$. This trend is likely linked to long-tailed deviations in the $StD$ trace plot for pre-dng, suggesting occasional inefficiencies in posterior exploration, particularly at the extremes of the parameter space. Furthermore, the left tail of the posterior distribution contains small peaks corresponding to low mutation rates, reinforcing the possibility that poor mixing or ABC rejection artifacts contribute to this trend. No significant interaction between biasVar was observed in relation to this trend.

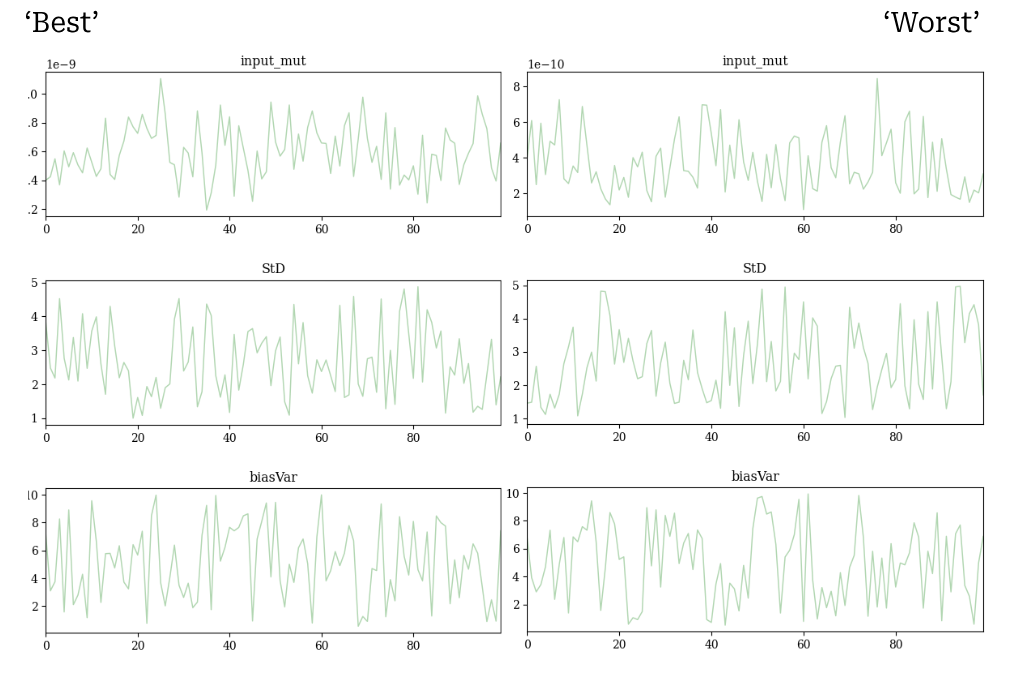

**Supplementary Fig. 2. MutSimABC Validation Trace Plots.** Trace plots for mutation rate (*μ*), elongation parameter ($StD$), and branching bias (*σ*) for the ‘best’ (left) and ‘worst’ (right) validation samples. Each sample contains exactly 100 accepted parameter sets. The best sample shows consistent mixing with high ESS values (67-100), indicating efficient posterior exploration. The worst sample exhibits slightly higher autocorrelation with lower ESS values (58-68), consistent with more challenging parameter space. Horizontal scatter indicates good mixing; clustering or trends would suggest poor sampling. Both cases demonstrate adequate posterior sampling given the limited number of accepted samples, with no evidence of non-convergence or stuck chains. *Trace Plots created with ArviZ [20].*

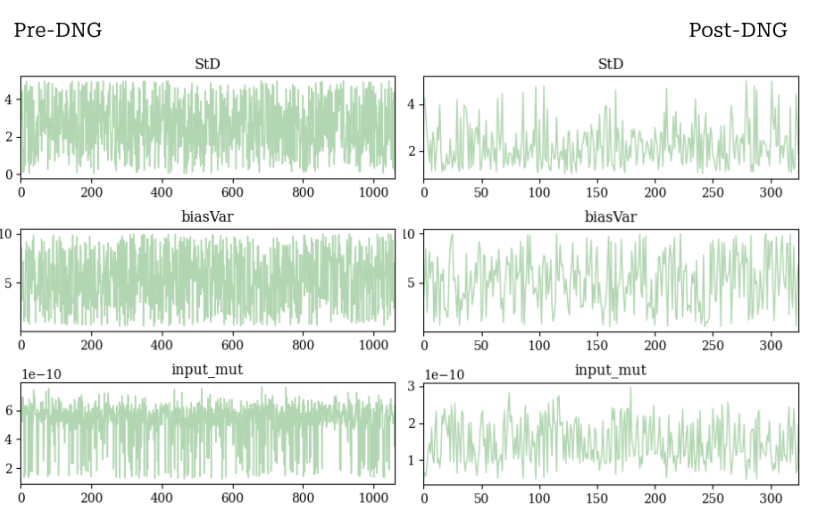

**Supplementary Fig. 3. MutSimABC Application to *E. melliodora* Trace Plots.** Trace plots for mutation rate (μ), elongation parameter (StD), and branching bias (σ) accepted samples for the ‘pre-DNG (left) and ‘post-DNG’ (right) empirical *E. melliodora* datasets -where DNG = topological filtering (Orr et al. (2020). The pre-DNG trace plot (left) shows some inconsistency in mixing, with long tails in certain regions indicating occasional inefficiencies in exploring the extremes of the posterior distribution. Specifically, the long tails likely correlate to the left tail and small peaks observed in the pre-DNG posterior. This observation aligns with the lower tail ESS (707) compared to the bulk ESS (970), indicating that the chain may struggle to sample the lower extremes of the mutation rate posterior fully. The post-DNG trace plot (right) shows relative consistency, with fewer streaks of similar values than parameters StD and biasVar. Bulk ESS (263) and tail ESS (283) indicate lower sampling efficiency compared to pre-DNG, which can be attributed to the smaller number of accepted samples - 350 out of 10,000 trials, compared to 1,020 accepted samples for pre-DNG. *Trace Plots created with ArviZ [20].*
